# Mind the Gap: Does Brain Age Improve Alzheimer’s Disease Prediction?

**DOI:** 10.1101/2024.11.16.623903

**Authors:** Trevor Wei Kiat Tan, Kim-Ngan Nguyen, Chen Zhang, Ru Kong, Susan F Cheng, Fang Ji, Joanna Su Xian Chong, Eddie Jun Yi Chong, Narayanaswamy Venketasubramanian, Csaba Orban, Michael W. L. Chee, Christopher Chen, Juan Helen Zhou, B. T. Thomas Yeo, the Alzheimer’s Disease Neuroimaging Initiative, the Australian Imaging Biomarkers and Lifestyle Study of Aging

## Abstract

Brain age is widely regarded as a powerful marker of *general* brain health. Brain age models are typically trained on large datasets to predict chronological age, which may offer advantages in predicting *specific* health outcomes, much like the success of finetuning large language models for specific applications. However, it is also well-accepted that machine learning models trained to directly predict specific outcomes (i.e., direct models) often outperform those trained on surrogate objectives. Therefore, despite their much larger training data, it is unclear whether brain age models outperform direct models in predicting specific brain health outcomes. Here, we compare large-scale brain age models (pretrained on 53,542 participants) and direct models for predicting specific health outcomes related to Alzheimer’s Disease (AD) dementia. Using anatomical T1 scans from three continents (N = 1,848), we find that summarizing brain age with a single scalar (i.e., brain age gap) led to poor prediction performance. Using higher-dimensional intermediate representations of brain age models led to better prediction, but was still worse than direct models without finetuning. Using intermediate representations of finetuned brain age models was necessary to achieve similar performance as direct models. Overall, our results do not discount brain age as a useful marker of general brain health, but suggest that using chronological age as a pretraining target might be suboptimal for predicting specific health outcomes.

## 1. INTRODUCTION

There is significant interest in using biological age as a marker of disease risk and mortality (Belsky et al., 2015; Chen et al., 2016; Tian et al., 2023). In the case of brain age, this involves training a machine learning model to predict chronological age from brain imaging data of healthy individuals (Dosenbach et al., 2010; Franke et al., 2010; Cole et al., 2017). The brain age gap (BAG) – the difference between predicted and chronological age – serves as a marker of accelerated aging and development. A positive BAG is associated with worse cognitive performance in older adults (Wrigglesworth et al., 2022; Cumplido-Mayoral et al., 2024), better cognitive performance in healthy children (Erus et al., 2014; Cheng et al., 2024), brain disorders (Kaufmann et al., 2019; Han et al., 2021; Constantinides et al., 2023), poor physical health (Franke et al., 2013; Ronan et al., 2016; Cole, 2020) and mortality (Cole et al., 2018; Paixao et al., 2020). Overall, these studies suggest the utility of BAG as a marker of *general* brain health.

In addition to associations at the group-level, BAG has been used to directly predict individual-level mortality (Cole et al., 2018), predict progression of mild cognitive impairment (MCI) to AD dementia (Gaser et al., 2013; Löwe et al., 2016; Choi et al., 2023) and classify psychiatric disorders (Koutsouleris et al., 2013; Leonardsen et al., 2022). However, summarizing a person’s brain health with a single number (BAG) might lose too much information. Therefore, some studies extract intermediate-level representations from pretrained brain age models, which are then used as input features for training new models to predict MCI progression (Gao et al., 2020), and classify neurological disorders (Leonardsen et al., 2022; Zheng et al., 2022). Finally, when deep neural networks (DNNs) are used for brain age prediction, the resulting models can be finetuned to diagnose brain disorders (Bashyam et al., 2020; Lu et al., 2022). The fine-tuning process can improve prediction by enabling the pretrained model to adapt to the unique characteristics of the new dataset – such as demographics or MRI scanner specifications – which may differ significantly from the data used to train the brain age model. Overall, these studies have demonstrated the utility of brain age models to predict *specific* health outcomes.

However, it remains unclear whether brain age derived models are better than models directly trained to predict specific health outcomes, which we refer to as “direct models”. On the one hand, there are orders of magnitude more brain imaging data with age-only information, compared with brain imaging data with target outcomes. Therefore, similar to the success of finetuning large language models for specific tasks (Yang et al., 2022; Tinn et al., 2023), a brain age model pretrained on tens of thousands of participants might yield better target prediction than direct models trained on hundreds of participants. On the other hand, a well-accepted machine learning principle is that training a model to directly predict a target variable of interest yields better prediction performance than training the model to predict a surrogate variable (that is only correlated with the target variable). Therefore, direct models might perform better than brain age derived models.

Motivated by this question, here we compare brain age derived models and direct models in two classification tasks. The first task is to predict whether a participant is cognitively normal (CN) or has AD dementia. We refer to this task as AD classification. The second task is to predict whether a participant with MCI would progress to AD dementia within 3 years. We refer to this task as MCI progression prediction. We chose these tasks because previous studies have suggested that brain age derived models can perform well in these tasks (Gaser et al., 2013; Bashyam et al., 2020; Lu et al., 2022). A previous study suggested that among nine brain disorders, T1 brain age gap between dementia and healthy controls was the largest with Cohen’s d of 1.03, compared with Cohen’s d of 0.07 to 0.74 for other brain disorders (Kaufmann et al., 2019). Given that brain age might be especially sensitive to dementia, the AD classification and MCI progression prediction tasks are natural benchmarks for brain age models.

Here, we consider the brain age model trained on one of the largest and most diverse datasets assembled (N = 53,542; Leonardsen et al., 2022). The Leonardsen brain age model utilizes the same convolutional neural network architecture as the winner of the Predictive Analysis Challenge for brain age prediction in 2019 (Gong et al., 2021; Peng et al., 2021). At the time of publication, the brain age model achieved state-of-the-art performance on data from unseen MRI scanners (Leonardsen et al., 2022). Intermediate representations from the pretrained model could also be used to classify various brain disorders via a transfer learning procedure (Leonardsen et al., 2022). As such, we believe the pretrained brain age model remains one of the best in the field. Evaluation was performed using anatomical T1 scans from three datasets (N = 1,848). We note that the evaluation datasets were not used to train the pretrained brain age model, so are truly out of sample.

Consistent with previous work (Leonardsen et al., 2022), we found that classifiers trained from intermediate representations of the pretrained brain age model (brainage64D) perform a lot better than BAG, suggesting that too much information is lost when summarizing a person’s biological age with a single number (i.e., BAG). Yet, direct models perform significantly better than brainage64D. Finetuning the brainage64D classifiers yields similar prediction performance to the direct models, but do not outperform direct models even when sample size is small (N = 50). Overall, our results do not dispute transfer learning as a general strategy to improve prediction or that brain age is a powerful marker of general brain health. However, our results highlight the misalignment between chronological age as a surrogate prediction target and the objective of predicting AD-related outcomes.

## 2. METHODS

### 2.1. Datasets

In this study, we considered three datasets: the Alzheimer’s Disease Neuroimaging Initiative (ADNI) dataset (Mueller et al., 2005; Jack et al., 2008; Jack et al., 2010), the Australian Imaging, Biomarkers and Lifestyle (AIBL) study (Ellis et al., 2009; Ellis et al., 2010; Fowler et al., 2021) and the Singapore Memory Aging and Cognition center (MACC) Harmonization cohort (Hilal et al., 2015; Xu et al., 2015; Chong et al., 2017; Hilal et al., 2020). Each dataset included both MRI data and clinical data collected at multiple timepoints.

In the first classification task, our goal was to predict whether an individual was cognitively normal (CN) or diagnosed with AD dementia at baseline using the baseline anatomical T1 scan. We refer to this task as AD classification. In the second classification task, our goal was to predict whether an individual with mild cognitive impairment (MCI) at baseline progressed to AD dementia within 36 months (i.e., progressive MCI or pMCI) or remained mild cognitively impaired (i.e., stable MCI or sMCI) based on the baseline anatomical T1 scan. We refer to this task as MCI progression prediction. Individuals who (1) exhibited more than one diagnosis changes (e.g., MCI → AD → MCI), (2) reverted to CN (i.e., MCI → CN), or (3) had missing diagnoses (such that we could not determine whether the individual should be considered pMCI or sMCI) were excluded. We note that there were no overlapping participants used for AD classification and MCI progression prediction tasks.

For the ADNI dataset, we considered participants from ADNI-1, ADNI-Go/2, and ADNI-3. T1 scans were acquired using 1.5T and 3T scanners from Siemens, Philips, and General Electric. At baseline, there were 2,039 scans, comprising 942 CN individuals, 432 individuals with AD dementia, 391 individuals who were sMCI, and 274 individuals who were pMCI. For the AIBL dataset, T1 scans were collected from Siemens 3T scanners. At baseline, there were 580 scans from 479 CN individuals, 78 individuals with AD dementia, 13 individuals who were sMCI, and 10 individuals who were pMCI. For the MACC dataset, T1 scans were collected from a Siemens 3T Tim Trio scanner, and a Siemens 3T Prisma scanner. At baseline, there were 457 scans from 132 CN individuals, 207 individuals with AD dementia, 79 individuals who were sMCI, and 39 individuals who were pMCI.

### 2.2. Sample stratification

Following Leonardsen and colleagues (Leonardsen et al., 2022), for the AD classification task, age and sex matching were performed for each scanner model to ensure the same age and sex distribution between the two diagnostic groups. For instance, suppose Scanner Model X in Dataset A has more participants with AD dementia than CN participant, then for each CN participant, we found the closest matching AD participant in terms of sex and age. Once all the CN participants were successfully matched, any excess AD participants were excluded from subsequent analyses. After matching, there were 636 CN participants and 636 participants with AD dementia. Figure 1A and Table 1 show the age and sex distributions of participants before and after matching. The same matching procedure was performed for the MCI progression prediction task (Figure 1B and Table 2), yielding 288 participants with sMCI and 288 participants with pMCI.

**Figure 1.**
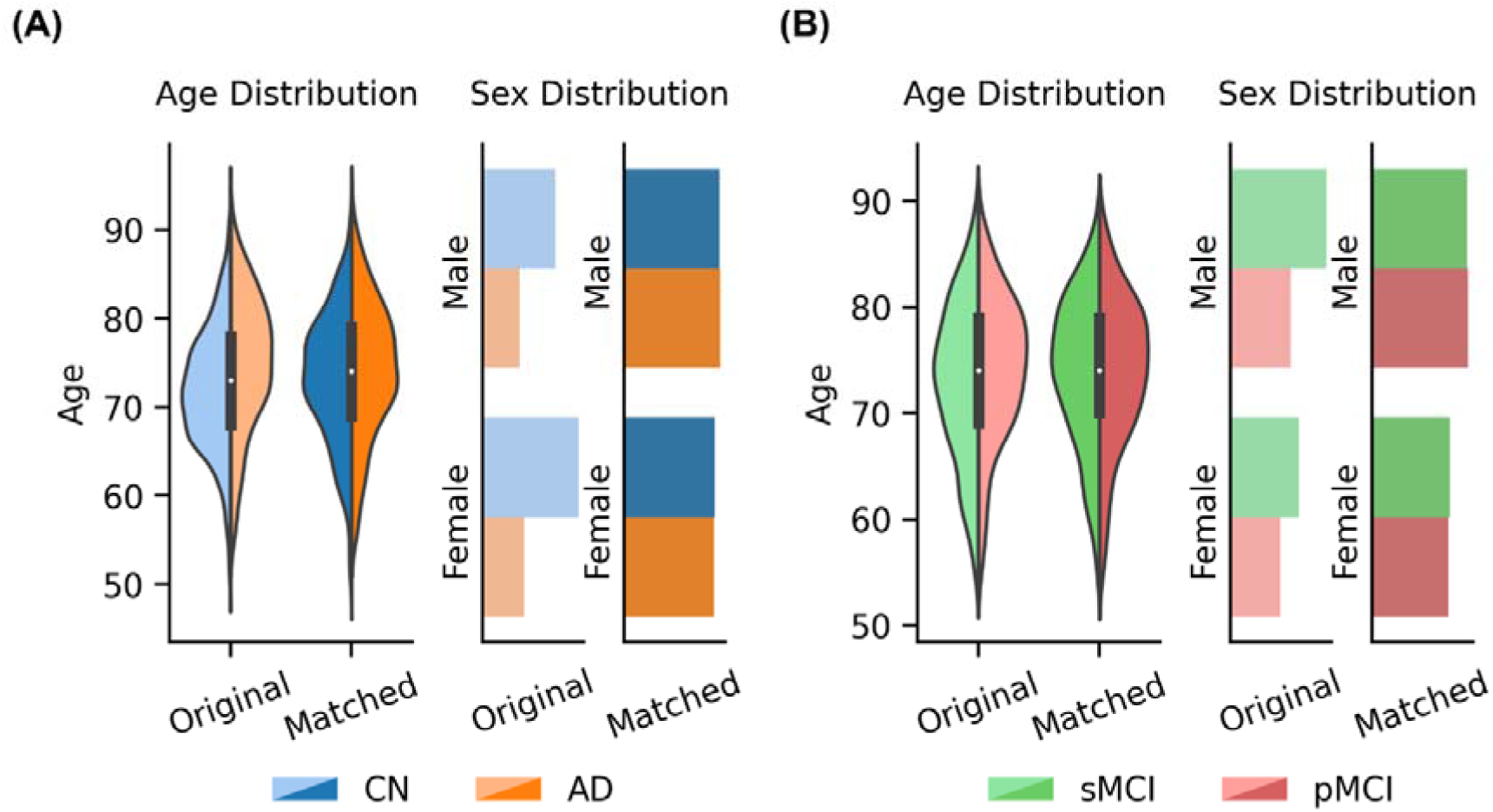
Age and sex distributions for each group before and after matching. (A) Age and sex distributions of CN participants and participants with AD dementia, before and after matching. (B) Age and sex distributions of participants with sMCI and pMCI, before and after matching.

**Table 1.**
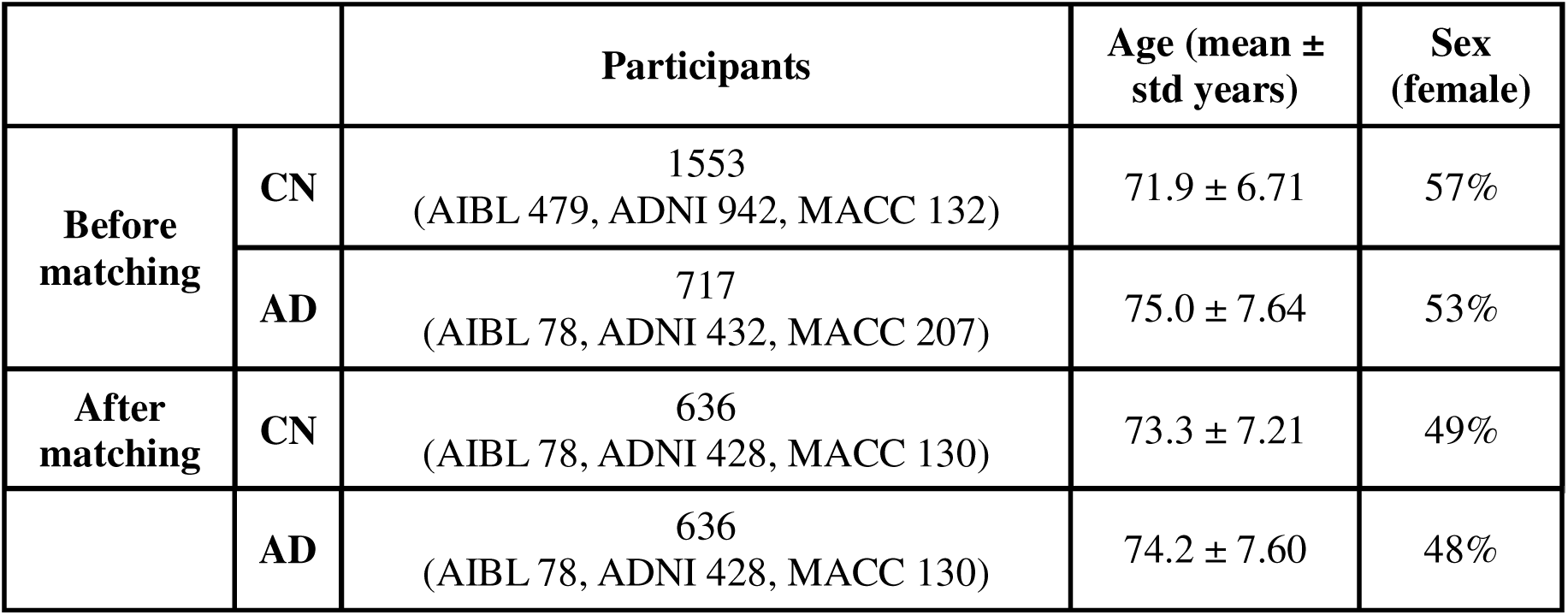
Participant demographics of three datasets (AIBL, ADNI, MACC) used for AD classification before and after matching age and sex. After matching, the paired t-test p value between the age of CN participants and participants with AD dementia was 0.031. For sex, the p value for the chi-square goodness of fit test between the two diagnostic groups was 0.99.

**Table 2.** Participant demographics of three datasets (AIBL, ADNI, MACC) used for MCI progression prediction before and after matching age and sex. After matching, the paired t-test p value between the age of the two groups (sMCI and pMCI) was 0.87. For sex, p value for chi-square goodness of fit test between the two groups was 0.99.

|  |  | **Participants** | **Age (mean years ± SD)** | **Sex (female)** |
| --- | --- | --- | --- | --- |
| **Before matching** | **sMCI** | 483<br>(AIBL: 13, ADNI 391, MACC: 79) | 73.0 ± 7.38 | 42% |
| | **pMCI** | 323<br>(AIBL: 10, ADNI 274, MACC: 39) | 74.2 ± 6.83 | 45% |
| **After matching** | **sMCI** | 288<br>(AIBL 10, ADNI 239, MACC 39) | 73.9 ± 6.93 | 45% |
| | **pMCI** | 288<br>(AIBL 10, ADNI 239, MACC 39) | 74.0 ± 6.87 | 44% |

### 2.3. Train, validation, and test split

For the AD classification task, we split the matched participants into a development set and test set, maintaining an 80:20 ratio. The development set was further divided into a training set and a validation set, also maintaining an 80:20 ratio. This results in training-validation-test split ratio of 64:16:20. The split into training, validation and test sets was performed separately for each scanner model of each dataset.

In general, the training set was used to train the parameters of each classification model. The validation set was used for early stopping (see details in Section 2.6). Finally, the test set was used to evaluate the performance of the model. This procedure was repeated 50 times with a different split of the participants into training, validation and test sets. The same procedure was repeated for the MCI progression prediction task.

### 2.4. T1 preprocessing

We performed the same T1 preprocessing as Leonardsen et al., 2022. Briefly, the T1 scans underwent skull stripping using FreeSurfer recon-all (Ségonne et al., 2004; Reuter et al., 2010), which generated a brain mask to remove non-brain areas and the skull. Subsequently, the brain was aligned to the standard FSL orientation via fslreorient2std (Jenkinson et al., 2012). Afterward, the images were linearly registered to MNI152 space using FLIRT (Jenkinson & Smith, 2001; Jenkinson et al., 2002; Greve & Fischl, 2009) with linear interpolation and a rigid body transformation with 6 degrees of freedom. The registration process used the FSL MNI152 1mm template. After registration, the images were cropped along the borders at [6:173, 2:214, 0:160] (using python indexing convention), resulting in 3D volumes of dimensions 167 × 212 × 160. This cropping procedure resulted in a compact cuboid that preserved almost all brain-related information. Finally, the voxel intensity values of all images were normalized to a range of [0, 1] by dividing all voxel intensities by 255.

### 2.5. Neural network backbone

The pretrained brain age model (Leonardsen et al., 2022) utilized the Simple Fully Convolutional Network (SFCN) backbone (Peng et al., 2021; Leonardsen et al., 2022). Therefore, we used the SFCN architecture for all tested models (Figure 2). The SFCN backbone comprised 5 repeated convolutional blocks and an appended 6^th^ convolutional block. Each of the 5 repeated convolutional blocks consisted of a 3D convolutional layer with a filter size of (3, 3, 3), zero padding with a padding size of 1, and a stride of 1. This results in convolutional layers of the same size as the input image. Each convolutional layer is followed by a batch normalization layer, rectified linear activation function (ReLU) activation, and a max pooling layer with a pooling size of (2, 2, 2). This results in the output size of each convolutional block being reduced by half across the height, width, and depth, compared to the input size. The appended 6^th^ convolutional block incorporates a channel-wise convolutional layer, a final batch normalization layer, and a global average pooling layer. The number of filters used in the convolutional layers are [32, 64, 128, 256, 256, 64], so the output of the global average pooling layer is of length 64. In the SFCN-regression model (Leonardsen et al., 2022), the 64 features were entered into a fully connected layer (with one output node) to directly predict chronological age. Because the pretrained SFCN-regression model achieved the best brain age prediction performance on external data (Leonardsen et al., 2022), we used the SFCN-regression model for the above analyses.

**Figure 2.**
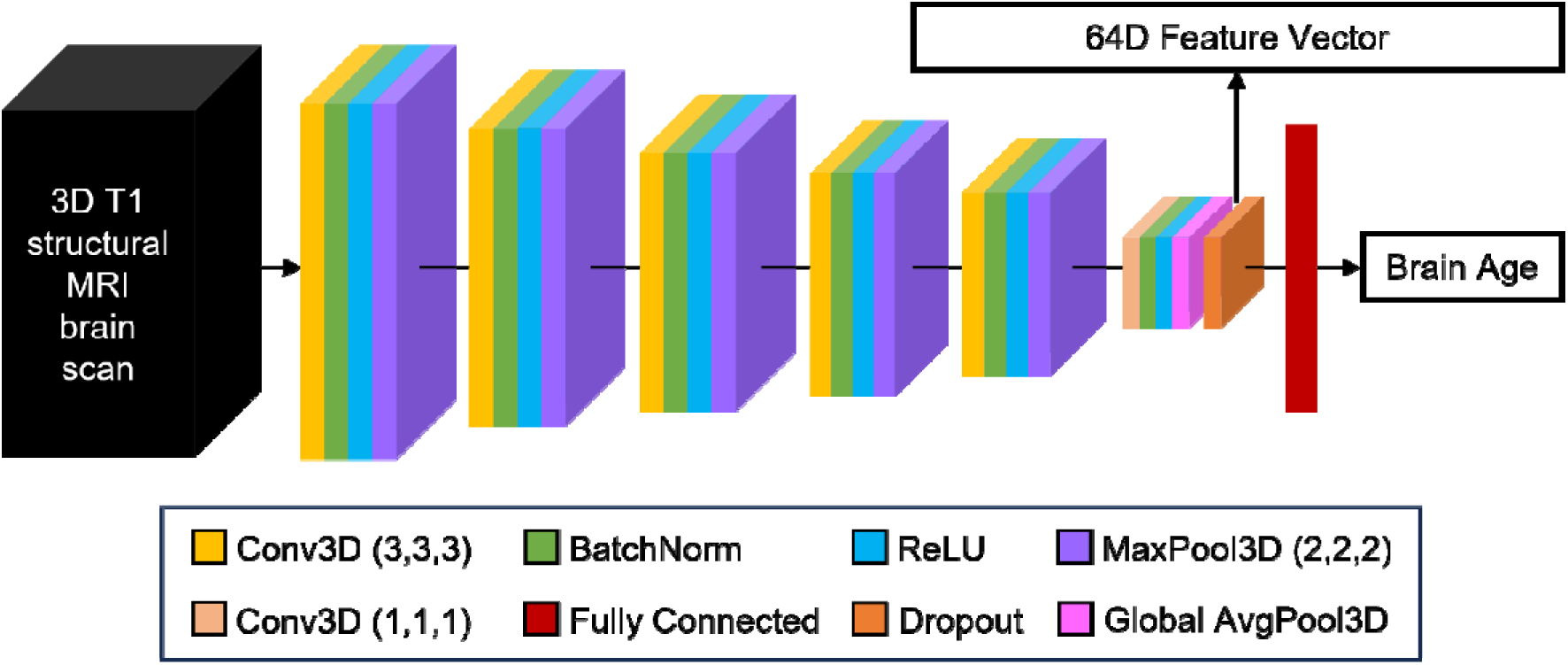
Schematic of Simply Fully Convolutional Network (SFCN) architecture (Peng et al., 2021; Leonardsen et al., 2022). The number of filters used in the convolutional layers were [32, 64, 128, 256, 256, 64]. The output of the global average pooling layer was of length 64. In the pretrained SFCN-regression model (Leonardsen et al., 2022), the 64-dimensional output of the global average pooling layer was fed into a fully connected layer (with one output node) to directly predict chronological age.

### 2.6. AD classification models

We compared five approaches for AD classification (Figure 3): Direct approach (Figure 3A), brain age gap (BAG; Figure 3B), BAG-finetune (Figure 3C), Brainage64D (Figure 3D) and Brainage64D-finetune (Figure 3E). All computations involving SFCN utilized NVIDIA RTX 3090 GPUs with 24GB memory and CUDA 11.0. Table 3 summarizes the amount of training/validation data and number of trainable parameters available to all models.

**Figure 3.**
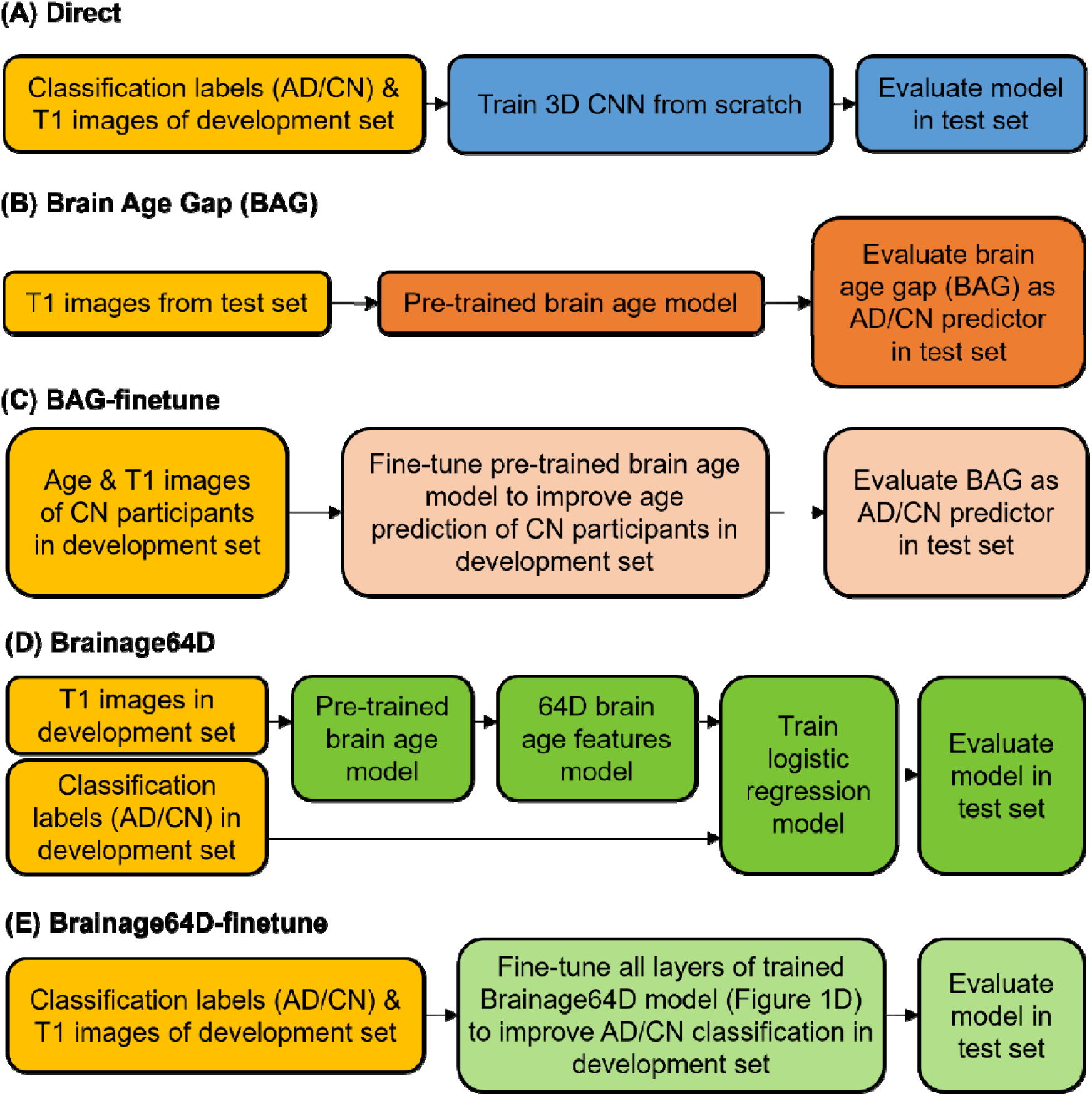
Workflow of five AD classification models. Given a T1 MRI scan, each model predicted a classification label (AD or CN). “Direct” is a model trained from scratch, while the other four models involved a brain age model that was previously trained from 53,542 participants across diverse datasets (Leonardsen et al., 2022). (A) “Direct” used the same 3D CNN architecture as the brain age model, except for the final binary classification layer. Classification labels (AD or CN) and T1 images of the development set were used to train the 3D CNN from scratch. (B) BAG model subtracted the chronological age of each test participant from the predicted age of the pretrained brain age model. The resulting brain age gap (BAG) was used as the AD/CN predictor. (C) Age & T1 images of CN participants in the development set were used to finetune the pretrained brain age model to improve age prediction of CN participants in development set. The finetuned brain age model was then used to compute BAG of each test participant, and the resulting BAG was used as the AD/CN predictor. (D) “Brainage64D” extracted 64-dimensional (64D) brain age features from the output of the global averaging pooling layer of the pretrained brain age model. The 64D features and classification labels (AD/CN) of the development set were used to train a logistic regression model. The final model consisted of the concatenation of the pretrained brain age model up to (and including) the global average pooling layer and the trained logistic regression model. (E) “Brainage64D-finetune” involved finetuning all layers of the trained Brainage64D model (from panel D) to improve AD classification in the development set.

**Table 3.** Comparison of sample sizes and number of trainable parameters across different models for AD classification. Compared with the direct model, we note that Brainage64D-finetune had the same training and validation set sizes, as well as the same number of trainable parameters, but also had a massive sample for chronological age pretraining.

|  | Brain age pretraining sample size | Training+validation set sizes | # trainable parameters |
| --- | --- | --- | --- |
| Direct | 0 | 997 | 2,950,466 |
| BAG | 53,542 | 0 | 0 |
| BAG-finetune | 53,542 | 499 | 2,950,401 |
| Brainage64D | 53,542 | 997 | 65 |
| Brainage64D-finetune | 53,542 | 997 | 2,950,466 |

#### 2.6.1. Direct approach

In the “Direct” approach, we trained a SFCN model from scratch to predict the classification label. More specifically, we replaced the fully connected layer (with one output node) of the SFCN-regression model with a fully connected layer with two output nodes for CN and AD classification labels respectively (Figure 3A). Stochastic gradient descent (SGD) was used to minimize the cross-entropy loss. Because of the computational cost, no hyperparameter was tuned. Instead, we used a fixed set of hyperparameters: weight decay = 1e-4, dropout rate = 0.5, and initial learning rate = 0.1. The learning rate was decreased by a factor of 10 every 30 epochs. The training batch size was set to 6 due to GPU memory constraints. For each training-validation-test split, the Direct approach was trained from scratch (using randomly initialized weights) for 150 epochs in the training set. Model parameters with the highest area under the receiver operating characteristic curve (AUC) in the validation set was used for evaluation in the test set. The evaluation metric was also AUC. For each training-validation-test split, the training duration was approximately 5 hours.

#### 2.6.2. Brain age gap (BAG) approach

The remaining four approaches involved the pretrained SFCN-regression model (Leonardsen et al., 2022). The first brain-age approach was simply the brain age gap (BAG) generated by the pretrained SFCN-regression model (Figure 3B). More specifically, the preprocessed T1 image of a test participant was fed into the pretrained brain age model. The BAG was then defined as the predicted age minus the actual chronological age of the participant. If the BAG was above a certain BAG threshold, we would classify the participant as having AD dementia. Otherwise, we classified the participant as CN. By varying the BAG threshold, we could compute the area under the receiver operating characteristic curve (AUC) for this approach. We note that the training and validation sets were not used at all for the BAG approach, but were used for other brain age models (Sections 2.6.3 to 2.6.5).

#### 2.6.3. Brain age gap finetune (BAG-finetune) approach

As will be seen in Section 3.2, the BAG approach did not result in a good classification accuracy. One potential reason is that the new datasets (ADNI, AIBL and MACC) were too different from the original multi-site datasets used to train the brain age model (Leonardsen et al., 2022). Therefore, we considered the BAG-finetune approach (Figure 3C), in which the pretrained brain age model was finetuned to predict chronological age in the CN participants. More specifically, for each training-validation-test split, all weights of the pretrained SFCN-regression model were finetuned to improve the prediction of the chronological age of the CN participants in the training set using the mean absolute error (MAE) loss. The same hyperparameters were used as the Direct approach, except that the initial learning rate was set to a low value of 0.01 to avoid significant deviation from the original pretrained weights. The validation set was used to select the epoch with the hyperparameters that yielded the best chronological age prediction based on MAE. The finetuned brain age model was then used to generate brain age gap in each test participant, which was in turn used to compute AUC in the test set.

Some studies have suggested the need to calibrate brain age models. One common approach is posthoc calibration, which adjusts model predictions without altering the model itself, by applying a constant offset (Dular & Špiclin, 2024) or fitting a linear/nonlinear correction (Le et al., 2018; Beheshti et al., 2019; Smith et al., 2019). While such methods are computationally efficient, they do not modify the internal representations learned by the model.

In contrast, we employed a transfer learning-based calibration (Holderrieth et al., 2023; Cheng et al., 2024; Wood et al., 2024) for BAG-finetune (current section), where all weights in the brain age model are finetuned. This calibration improves chronological age prediction, reducing MAE from 4.4 to 3.5 in CN participants from the test set (of the AD-CN classification task). This transfer learning approach was also used for the “Brainage64D-finetune” (Section 2.6.5) and “Brainage64D-finetune-AD2prog” (Section 2.7.3) models, which provided a unified calibration method across the different brain age pipelines examined in our study.

Furthermore, Leonardsen and colleagues, who developed the brain age model used in the current study, asserted that “[f]rom a practical point of view this suggests that our trained model may be employed in other applications in new datasets, without the need for retraining or applying corrective procedures.” (Leonardsen et al., 2022). Consequently, we believe that it is reasonable to include BAG (Section 2.6.2) as an uncalibrated reference, in line with its original design (Leonardsen et al., 2022). Importantly, we also included calibrated brain age models in our comparisons, ensuring a fair and comprehensive evaluation of brain age and direct models.

#### 2.6.4. Brain age 64D (Brainage64D) approach

As will be seen, the BAG-finetune approach also did not perform well, so another possible hypothesis is that summarizing a participant with just a single scalar (brain age gap) might be losing too much information. Therefore, we considered the brain age 64D (Brainage64D) approach, in which a logistic ridge regression model was trained on the 64-dimensional output of the global averaging pooling layer in the pretrained SFCN-regression model (Figure 3D). The scikit-learn package (Pedregosa et al., 2011) was used. The logistic ridge regression model included an inverse regularization parameter λ (larger value indicated lower regularization). Model fitting was performed on the training set, and the inverse regularization parameter was determined based on AUC in the validation set.

The optimal hyperparameter was selected from 0.001, 0.01, 0.1, 1, 10, 100 or 1000. The best hyperparameter was then used to retrain the logistic regression model on the full development (training and validation) set. The final trained Brainage64D model was the concatenation of the pretrained SFCN-regression model up to (and including) the global average pooling layer and the trained logistic regression model. This final model was then evaluated in the test set.

### 2.6.5. Brain age 64D finetune (Brainage64D-finetune) approach

As will be seen in Section 3.2, the Brainage64D approach performed better than BAG (Section 2.6.2) and BAG-finetune (Section 2.6.3), but was still worse than the Direct approach. One potential reason is that the new datasets (ADNI, AIBL and MACC) were too different from the original multi-site datasets used to train the brain age model (Leonardsen et al., 2022).

Therefore, we considered a Brainage64D-finetune approach (Figure 3E), in which we finetuned the previously trained Brainage64D model (Section 2.6.4). All layers of the trained Brainage64D model were finetuned, giving Brainage64D-finetune an identical number of trainable parameters as the Direct approach (in Section 2.6.1). The same cost function and hyperparameters as the Direct approach were used, except that the initial learning rate was set to a low value of 0.01 to avoid significant deviation from the original pretrained weights. Model parameters with the highest AUC in the validation set was then used for evaluation in the test set.

### 2.7. MCI progression prediction models

We compared three approaches for MCI progression prediction (Figure 4): Direct approach (Figure 4A), Direct-AD2prog (Figure 4B) and Brainage64D-AD2prog (Figure 4C). All computations involving SFCN utilized NVIDIA RTX 3090 GPUs with 24GB memory and CUDA 11.0. Table 4 summarizes the amount of training/validation data and number of trainable parameters available to all models.

**Figure 4.**
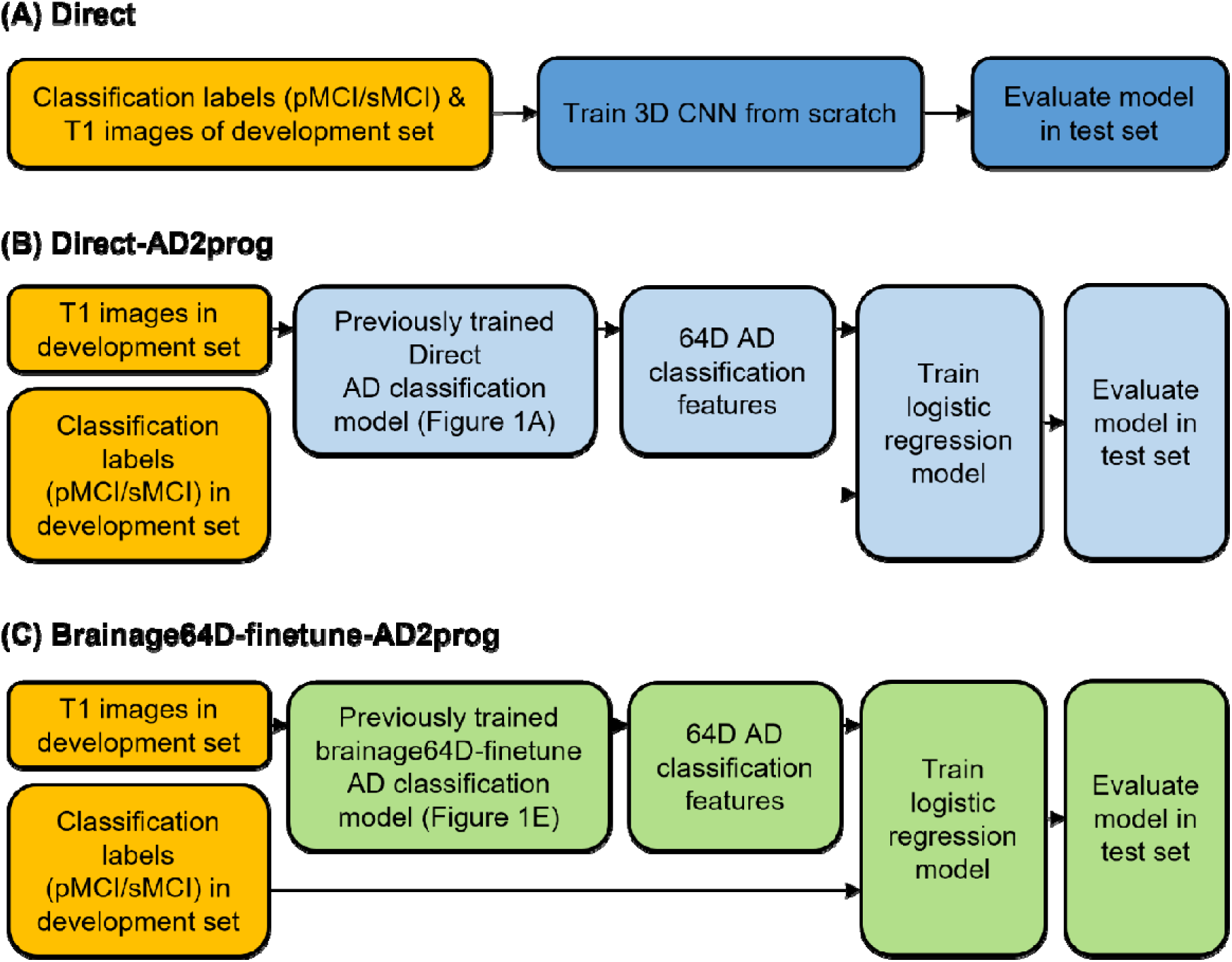
Workflow of three MCI progression prediction models. Given a T1 MRI scan, each model predicted a classification label (sMCI or pMCI). (A) “Direct” used the same 3D CNN architecture as the pretrained brain age model (Leonardsen et al., 2022), except for the final binary classification layer. Classification labels (sMCI or pMCI) and T1 images of the development set were used to train the 3D CNN from scratch. (B) “Direct-AD2prog” extracted 64-dimensional (64D) features from the output of the global averaging pooling layer of the previously trained Direct AD classification model (Figure 3A). The 64D features and classification labels (sMCI/pMCI) of the development set were used to train a logistic regression model. (C) “Brainage64D-finetune-AD2prog” extracted 64-dimensional (64D) features from the output of the global averaging pooling layer of the previously trained brainage64D-finetune AD classification model (Figure 3E). The 64D features and classification labels (sMCI/pMCI) of the development set were used to train a logistic regression model.

**Table 4.** Comparison of sample sizes and number of trainable parameters across different models for MCI progression prediction. Compared with the Direct-AD2prog model, we note that Brainage64D-finetune-AD2prog had the same training and validation set sizes, same AD classification sample size and same number of trainable parameters, but also had a massive sample for chronological age pretraining.

|  | Brain age pretraining sample size | AD classification pretraining sample size | training+validation set sizes | # trainable parameters |
| --- | --- | --- | --- | --- |
| Direct | 0 | 0 | 448 | 2,950,466 |
| Direct-AD2prog | 0 | 997 | 448 | 65 |
| Brainage64D-finetune-AD2prog | 53,542 | 997 | 448 | 65 |

#### 2.7.1. Direct approach

The “Direct” approach for predicting MCI progression is the same as “Direct” approach for AD classification. In other words, we trained a SFCN model from scratch to predict whether a participant was pMCI or sMCI. Similar to Section 2.6.1, we replaced the fully connected layer (with one output node) of the SFCN-regression model with a fully connected layer with two output nodes for sMCI and pMCI classification labels respectively (Figure 4A). Stochastic gradient descent (SGD) was used to minimize the cross-entropy loss. Because of the computational cost, no hyperparameter was tuned. Instead, we used a fixed set of hyperparameters: weight decay = 1e-4, dropout rate = 0.5, and initial learning rate = 0.1. The learning rate was decreased by a factor of 10 every 30 epochs. The training batch size was set to 6 due to GPU memory constraints. For each training-validation-test split, the Direct approach was trained from scratch (using randomly initialized weights) for 150 epochs in the training set. Model parameters with the highest area under the receiver operating characteristic curve (AUC) in the validation set was used for evaluation in the test set.

#### 2.7.2. Direct-AD2prog

As seen in Section 3.4, the prediction performance of the “Direct” approach was not good. Previous studies have suggested that adapting an AD classification task to predict MCI progression can improve prediction performance, compared with training a model from scratch (Oh et al., 2019; Lian et al., 2020; Wen et al., 2020). Therefore, we extracted the 64-dimensional output of the global averaging pooling layer of the previously trained Direct AD classification models (Section 2.6.1; Figure 3A). We then trained a logistic ridge regression model using the 64-dimensional features to predict whether a participant progressed to AD dementia. We refer to this approach as Direct-AD2prog (Figure 4B), where “AD2prog” refers to the fact that we transferred features from the AD classification model to build a new model to predict disease progression.

Consistent with Section 2.6.4, the logistic ridge regression model included an inverse regularization parameter λ (larger value indicated lower regularization), which was determined based on AUC in the validation set. The optimal hyperparameter was selected from 0.001, 0.01, 0.1, 1, 10, 100 or 1000. The best hyperparameter was then used to retrain the logistic regression model on the development (training and validation) set. The trained model was then evaluated on the test set.

#### 2.7.3. Brainage64D-finetune-AD2prog

Similar to Direct-AD2prog, we also extracted the 64-dimensional output of the global averaging pooling layer of the previously trained Brainage64D-finetune AD classification models (Section 2.6.5). We then trained a logistic ridge regression model using the 64-dimensional features to predict whether a participant progressed to AD dementia. We refer to this approach as Brainage64D-finetune-AD2prog (Figure 4C). Consistent with previous sections, we again select the optimal regularization hyperparameter (from 0.001, 0.01, 0.1, 1, 10, 100 or 1000) based on the validation set. The best hyperparameter was then used to retrain the logistic regression model on the full development (training and validation) set. The trained model was then evaluated on the test set. Overall, we note that Brainage64D-finetune-AD2prog has the same number of trainable parameters as Direct-AD2prog (Section 2.7.2).

### 2.8. Vary training and validation set sizes

Adapting a pretrained model (trained from large datasets) for a new classification task might be more advantageous when the sample size available for the new task is small. Therefore, we repeated the previous AD classification and MCI progression prediction tasks by varying the size of the development (training and validation) set.

Recall that we have previously repeated the training-validation-test procedure 50 times (Section 2.3). In the current analysis, for each of these 50 repetitions, we in turn randomly sub-sample the development (training and validation) set. In the case of AD classification, we varied the development set size as follows: 50, 100, 200, 300, 400, 500, 600, 700, 800, 900, and the maximum development set size of 997. In the case of MCI progression prediction, we varied the development set size as follows: 50, 100, 150, 200, 250, 300, 350, 400, and the maximum development set size of 448.

Consistent with previous analyses, the development set was in turn divided into training and validation sets with an 80:20 ratio. The test set was also maintained to be the same across all development set sample sizes, so that the prediction accuracies were comparable across development sample sizes.

In the case of AD classification, care was taken so that when the subsampled development set had the same numbers of CN participants and participants with AD dementia. Since BAG and BAG-finetune performed poorly in the main analysis (Figure 5), for this analysis we only considered Brainage64D, Brainage64D-finetune and Direct approach.

**Figure 5.**
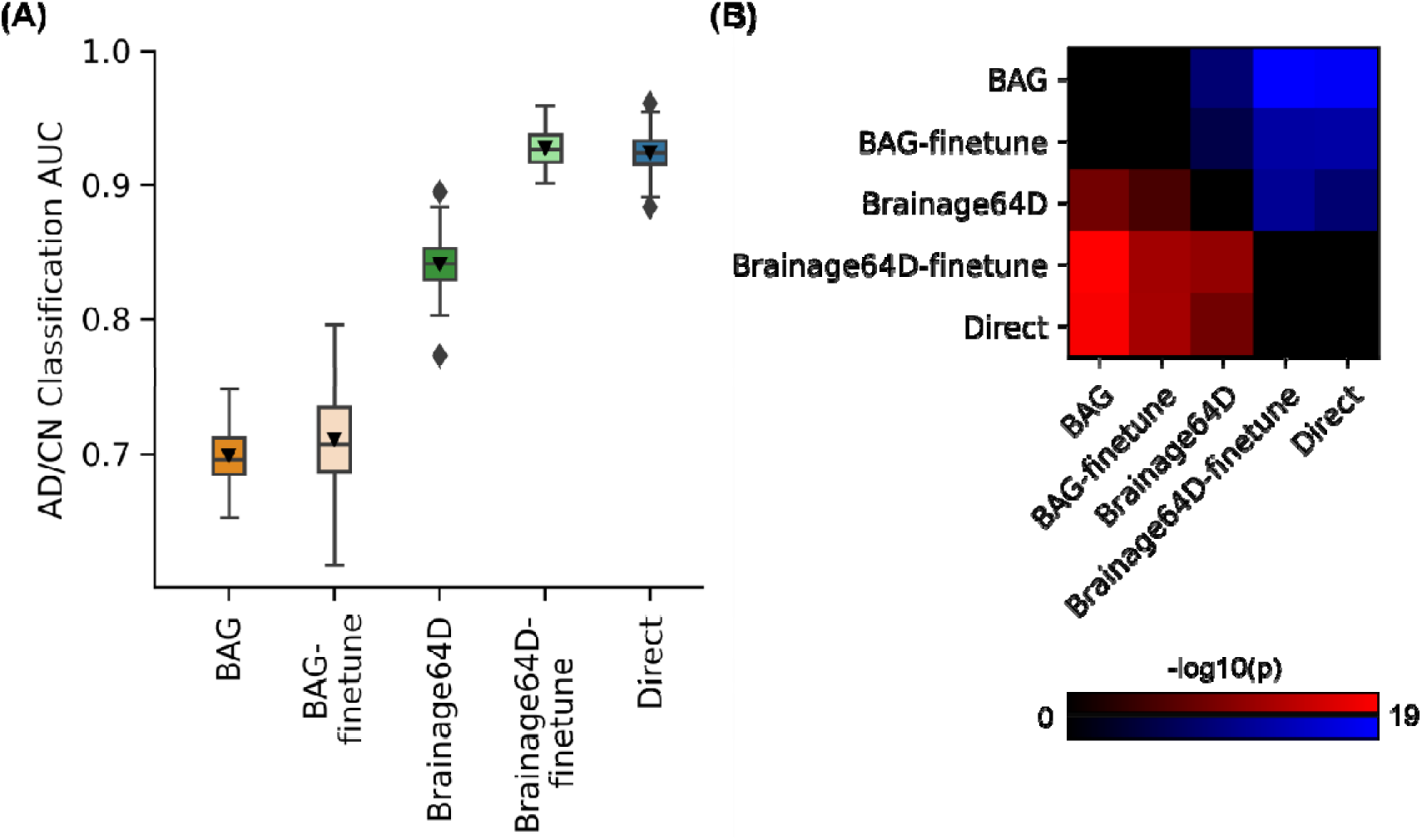
AD classification AUC. (A) Box plots show the test AUC across 50 random training-validation-test splits. For each box plot, the horizontal line indicates the median across 50 test AUC values. Triangle indicates the mean. The bottom and top edges of the box indicate the 25th and 75th percentiles, respectively. Outliers are defined as data points beyond 1.5 times the interquartile range. The whiskers extend to the most extreme data points not considered outliers. (B) Minus log10 p values between all pairs of AD classification approaches based on the corrected resampled t-test (Nadeau & Bengio, 2003). Larger - log10(p) indicates greater statistical significance. Non-black color indicates significant differences after false discovery rate (FDR) correction (q < 0.05). For each pair of approaches, red (or blue) color indicates that the approach on the row outperformed (or underperformed) the approach on the column. For example, if we focus on the “Direct” row, the red color indicates that “Direct” statistically outperformed BAG, BAG-finetune and Brainage64D.

In the case of MCI progression prediction, care was taken so that when the subsampled development set had the same numbers of sMCI and pMCI participants. Since Direct performed poorly in the main analysis (Figure 7), for this analysis we only considered Direct-AD2prog and Brainage64D-finetune-AD2prog. Furthermore, while we varied the development set size, the input features for both approaches were extracted from AD classification models trained from the full sample size (Section 2.7).

## 3. RESULTS

### 3.1. Overview

Here we considered data from three datasets: the Alzheimer’s Disease Neuroimaging Initiative (ADNI) dataset (Mueller et al., 2005; Jack et al., 2008; Jack et al., 2010), the Australian Imaging, Biomarkers and Lifestyle (AIBL) study (Ellis et al., 2009; Ellis et al., 2010; Fowler et al., 2021) and the Singapore Memory Aging and Cognition center (MACC) Harmonization cohort (Hilal et al., 2015; Xu et al., 2015; Chong et al., 2017; Hilal et al., 2020). More details about the datasets and preprocessing can be found in Methods (Sections 2.1 and 2.4).

Following Leonardsen and colleagues (Leonardsen et al., 2022), we considered age-matched and sex-matched cognitively normal (CN) participants and participants diagnosed with AD dementia (N = 1272) for the AD classification task. We also considered age-matched and sex-matched MCI participants who progressed to AD dementia within three years (i.e., progressive MCI or pMCI) and remained as MCI (i.e., stable MCI or sMCI) for the MCI progression task (N = 576). For more details, see Methods (Section 2.2).

For both classification tasks, we utilized a nested (inner-loop) cross-validation procedure, in which participants were assigned to a development set and a test set with an 80:20 ratio. The development set was in turn divided into a training set and a validation set with an 80:20 ratio. In general, models were trained on the training set and hyperparameters were tuned on the validation set. The final model was evaluated in the test set. This training-validation-test procedure was repeated 50 times for robustness. For more details, see Methods (Section 2.3).

### 3.2. Leveraging a pretrained brain age model does not improve AD classification over training a model directly

For AD classification, we compared five approaches (Figure 3) - the direct model and four brain age models (BAG, BAG-finetune, Brainage64D and Brainage64D-finetune) derived from a state-of-the-art pretrained brain age model (Leonardsen et al., 2022). The brain age model was previously trained on 53,542 participants across diverse datasets (Leonardsen et al., 2022). For more details, see Methods (Sections 2.5 and 2.6). Figure 5A shows the AD classification AUC (across 50 training-validation-test splits) for all five approaches. Figure 5B illustrates the p values from comparing pairs of approaches using the corrected resampled t-test (Nadeau & Bengio, 2003). Table 5 reports the actual p values.

**Table 5.**
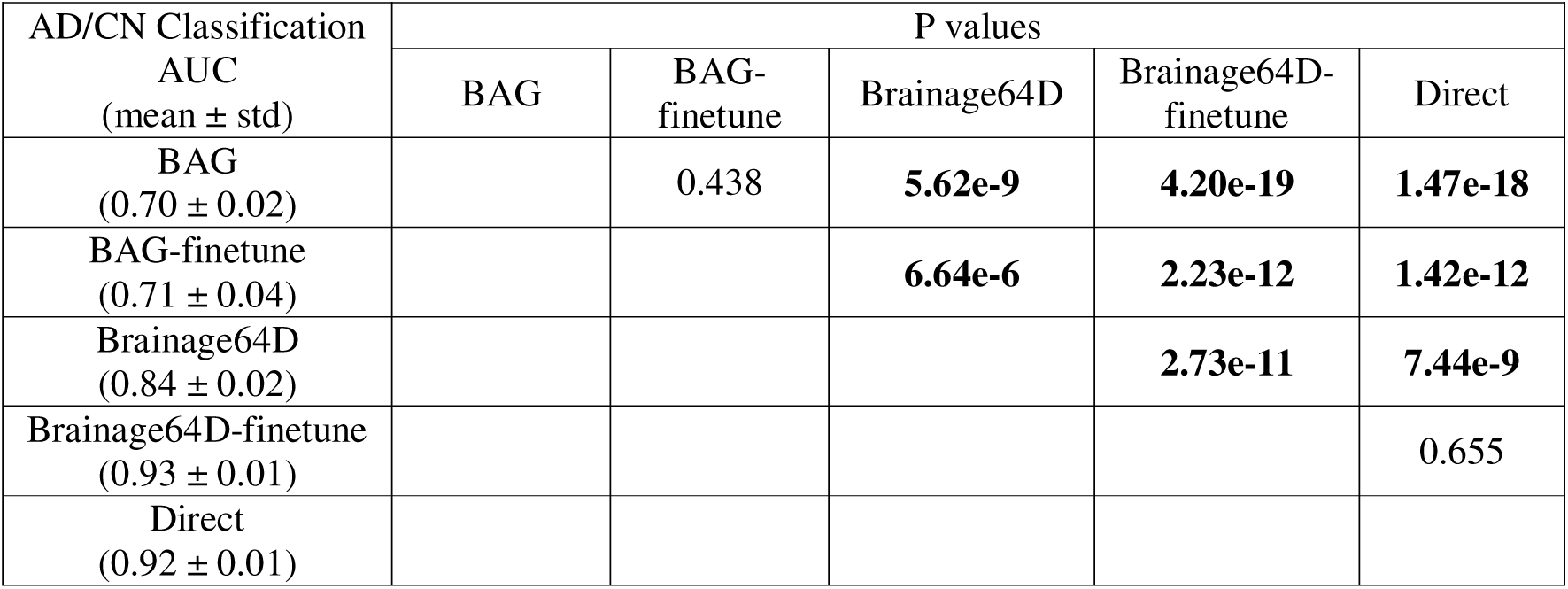
AD classification AUC (mean ± std) and uncorrected p values between pairs of approaches. AUC was averaged across 50 random training-validation-test splits. P values were computed using the corrected resampled t-test (Nadeau & Bengio, 2003). Bolded p values indicate statistical significance after FDR correction (q < 0.05).

| AD/CN Classification<br>AUC<br>(mean ± std) | P values | | | | |
| --- | --- | --- | --- | --- | --- |
|  | BAG | BAG-finetune | Brainage64D | Brainage64D-finetune | Direct |
| BAG<br>(0.70 ± 0.02) | | 0.438 | **5.62e-9** | **4.20e-19** | **1.47e-18** |
| BAG-finetune<br>(0.71 ± 0.04) | | | **6.64e-6** | **2.23e-12** | **1.42e-12** |
| Brainage64D<br>(0.84 ± 0.02) | | | | **2.73e-11** | **7.44e-9** |
| Brainage64D-finetune<br>(0.93 ± 0.01) | | | | | 0.655 |
| Direct<br>(0.92 ± 0.01) | | | | | |

Direct and Brainage64D-finetune performed the best, with no statistical difference between the two approaches. BAG and BAG-finetune performed the worst with no statistical difference between the two approaches. Brainage64D (without finetuning) achieved an intermediate level of performance. Importantly, the performance of Brainage64D (mean AUC=0.84) was comparable to the results reported by Leonardsen and colleagues (2022) (mean AUC=0.83).

Overall, this suggests that with the largest training set size of 997, leveraging a pretrained brain age model did not improve AD classification performance, compared with simply training a model from scratch.

### 3.3. Even when sample size is small, leveraging a pretrained brain age model does not improve AD classification over training a model directly

Adapting a pretrained model (trained from large datasets) for a new classification task might be more advantageous when the sample size available for the new task is small. Figure 6 shows the AD classification AUC (across 50 training-validation-test splits) for Direct, Brainage64D and Brainage64D-finetune across different development set sizes. Given their poor performance (Figure 5), we did not consider BAG and BAG-finetune in this analysis. Table 6 reports the actual AUC, while Table 7 reports the p values obtained from comparing the Direct approach with Brainage64D and Brainage64D-finetune using the corrected resampled t-test.

**Figure 6.**
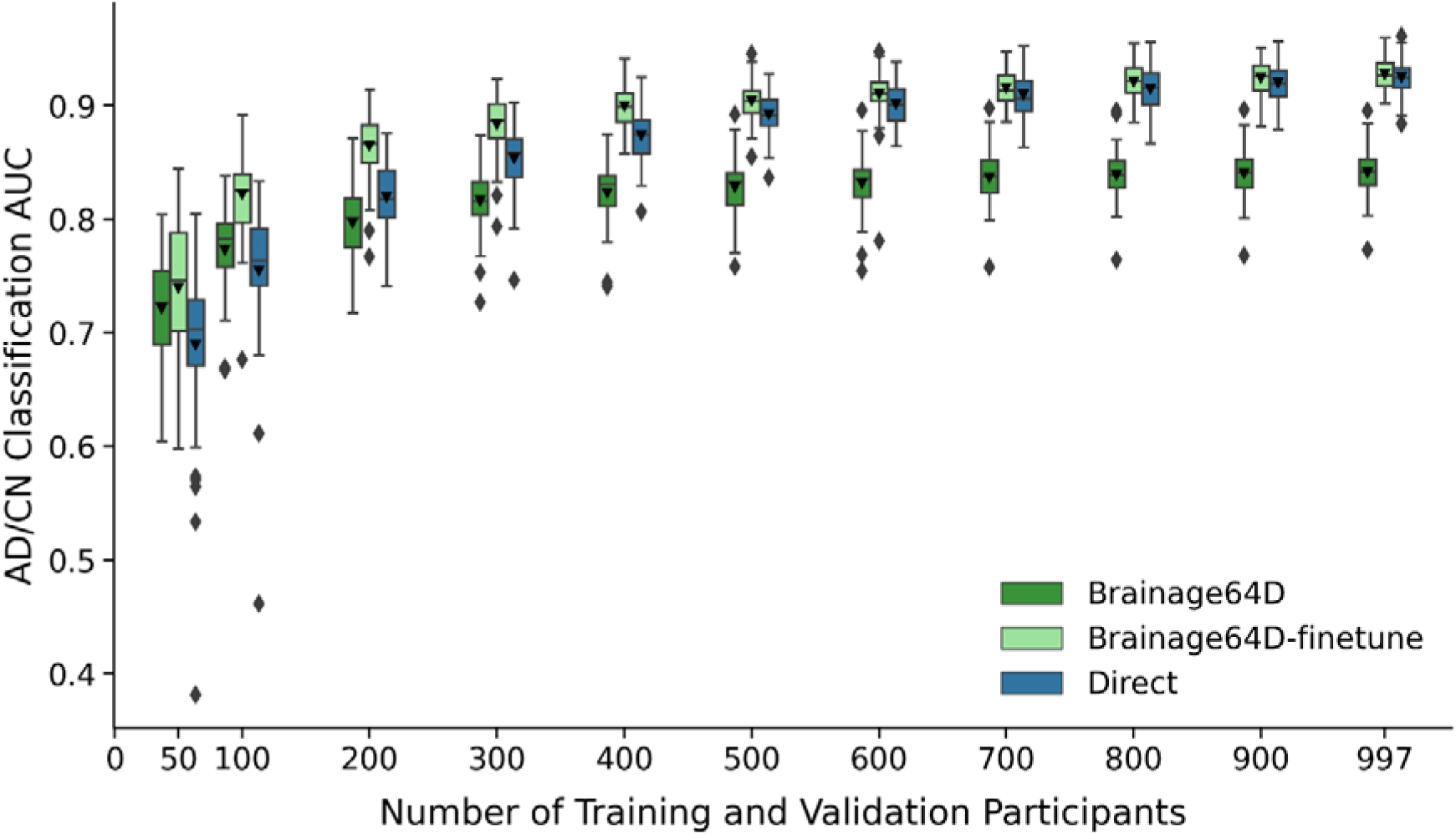
AD classification AUC of Brainage64D, Brainage64D-finetune, and Direct across different development set sizes. Boxplots showing test AUC across different development set sizes (number of training and validation participants). Test sets were identical across development set sizes, so AUCs were comparable across sample sizes. For each boxplot, the horizontal line indicates the median across 50 test AUC values. Triangle indicates the mean. The bottom and top edges of the box indicate the 25th and 75th percentiles, respectively. Outliers are defined as data points beyond 1.5 times the interquartile range. The whiskers extend to the most extreme data points not considered outliers.

**Table 6.** AD classification AUC (mean ± std) of different approaches for AD classification task across different development set sizes (i.e., number of training and validation participants). Given their poor performance (Figure 5), we did not consider BAG and BAG-finetune in this analysis.

| Number of training and validation participants | Brainage64D | Brainage64D-finetune | Direct |
| --- | --- | --- | --- |
| 50 | $0.7214 \pm 0.0465$ | $0.7393 \pm 0.0595$ | $0.6893 \pm 0.0742$ |
| 100 | $0.7729 \pm 0.0384$ | $0.8216 \pm 0.0368$ | $0.7547 \pm 0.0596$ |
| 200 | $0.7968 \pm 0.0319$ | $0.8643 \pm 0.0284$ | $0.8195 \pm 0.0307$ |
| 300 | $0.8160 \pm 0.0269$ | $0.8833 \pm 0.0251$ | $0.8533 \pm 0.0288$ |
| 400 | $0.8231 \pm 0.0277$ | $0.8996 \pm 0.0185$ | $0.8736 \pm 0.0221$ |
| 500 | $0.8279 \pm 0.0255$ | $0.9042 \pm 0.0163$ | $0.8919 \pm 0.0185$ |
| 600 | $0.8315 \pm 0.0246$ | $0.9101 \pm 0.0235$ | $0.9013 \pm 0.0195$ |
| 700 | $0.8362 \pm 0.0234$ | $0.9157 \pm 0.0144$ | $0.9094 \pm 0.0191$ |
| 800 | $0.8389 \pm 0.0226$ | $0.9211 \pm 0.0152$ | $0.9148 \pm 0.0182$ |
| 900 | $0.8403 \pm 0.0213$ | $0.9241 \pm 0.0152$ | $0.9200 \pm 0.0172$ |
| 997 | $0.8411 \pm 0.0213$ | $0.9276 \pm 0.0144$ | $0.9246 \pm 0.0146$ |

**Table 7.** Uncorrected p values between AUC of Direct and Brainage64D, as well as between Direct and Brainage64D-finetune across different development set sizes (i.e., number of training and validation participants). Bolded p values indicate statistical significance after FDR correction (q < 0.05). Given their poor performance (Figure 5), we did not consider BAG and BAG-finetune in this analysis.

| Number of training and validation participants | Direct vs Brainage64D | Direct vs Brainage64D-finetune |
| --- | --- | --- |
| 50 | 0.8816 | 0.8314 |
| 100 | 0.8696 | 0.5990 |
| 200 | 0.6743 | 0.2287 |
| 300 | 0.2941 | 0.2888 |
| 400 | 0.0368 | 0.1035 |
| 500 | **0.0010** | 0.3727 |
| 600 | **0.0001** | 0.5728 |
| 700 | **2.7106e-5** | 0.5171 |
| 800 | **2.0102e-6** | 0.3845 |
| 900 | **4.0756e-8** | 0.5238 |
| 997 | **7.4415e-9** | 0.6551 |

Brainage64D was numerically better than the Direct approach for development set sizes of 50 and 100, but the improvement was not statistically significant. The Direct approach was numerically better than Brainage64D from a development set size of 200 onwards, which became statistically significant when development set size was at least 500.

On the other hand, the Brainage64D-finetune was numerically better than the direct approach for all sample sizes, but differences were not significant even when sample size was very small (N = 50). Overall, this suggests that leveraging a pretrained brain age model did not improve AD classification over training a model directly.

### 3.4. Leveraging a pretrained brain age model does not improve MCI progression prediction over training a model directly

For the MCI progression prediction, we compared two direct models (Direct and Direct-AD2prog) with the best brain age derived model (Brainage64D-finetune-AD2prog; Figure 4). Both Direct-AD2prog and Brainage64D-finetune-AD2prog utilized intermediate representations (features) from the best AD classification models (Direct and Brainage64D-finetune) as inputs to train a new classifier for predicting MCI progression (Figure 4). For more details, see Methods (Section 2.7).

Figure 7 shows the MCI progression prediction AUC for all three approaches. Table 8 reports the p values from comparing pairs of approaches using the corrected resampled t-test (Nadeau & Bengio, 2003). Both Direct-AD2prog and Brainage64D-finetune-AD2prog performed better than the direct approach. There was no statistical difference between Direct-AD2prog and Brainage64D-finetune-AD2prog.

**Figure 7.**
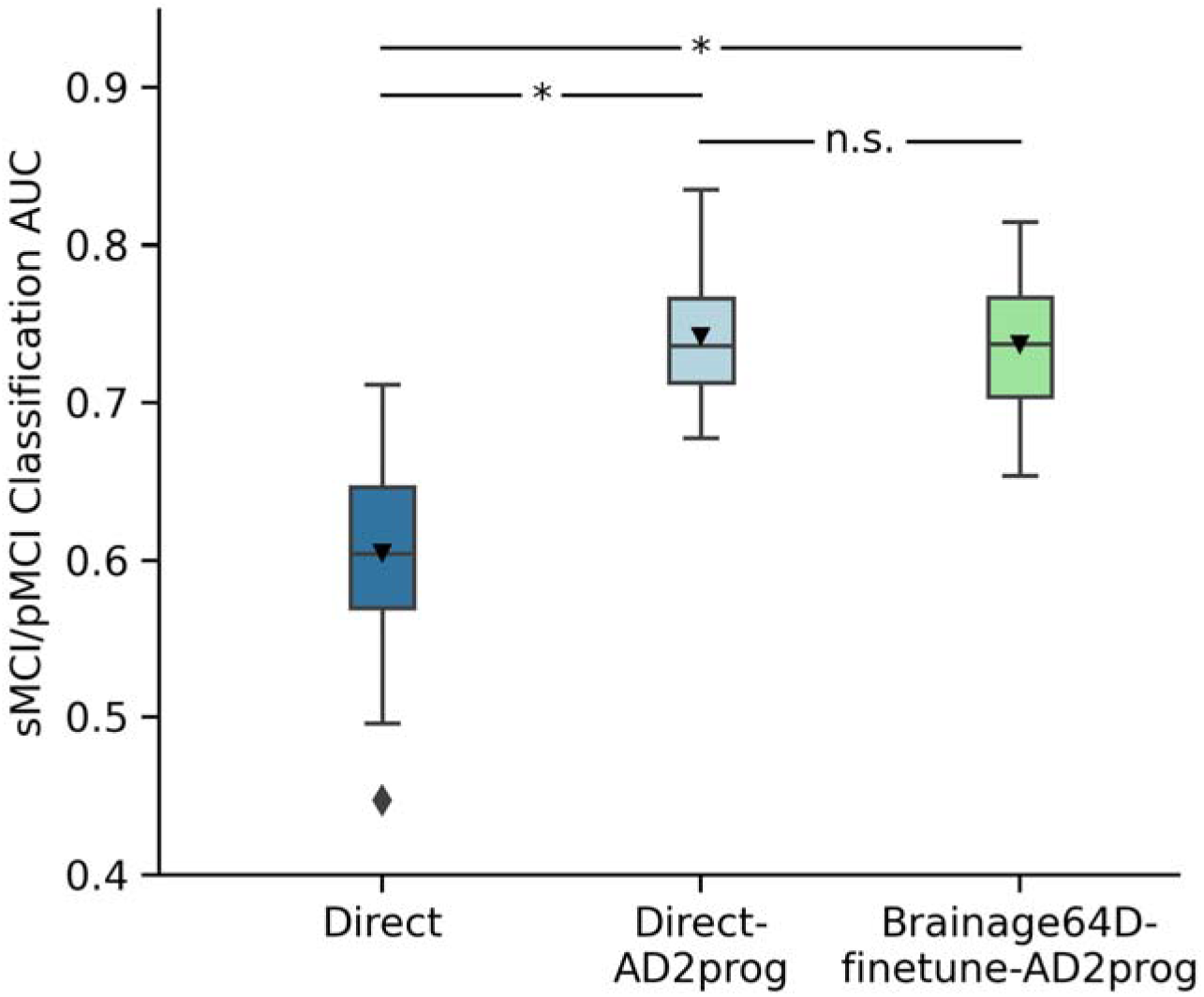
MCI progression prediction AUC. Box plots show the test AUC across 50 random training-validation-test splits. For each box plot, the horizontal line indicates the median across 50 test AUC values. Triangle indicates the mean. The bottom and top edges of the box indicate the 25th and 75th percentiles, respectively. Outliers are defined as data points beyond 1.5 times the interquartile range. The whiskers extend to the most extreme data points not considered outliers. P values are computed using the corrected resampled t-test. “*” indicates statistical significance after FDR correction (q < 0.05) and “n.s.” indicates not significant after FDR correction.

**Table 8.** MCI progression prediction AUC (mean ± std) and uncorrected p values between pairs of approaches. AUC was averaged across 50 random training-validation-test splits. P values were computed using the corrected resampled t-test (Nadeau & Bengio, 2003). Bolded p values indicate statistical significance after FDR correction (q < 0.05).

| sMCI/pMCI classification<br>AUC (mean ± std) | P values | | |
| --- | --- | --- | --- |
|  | Direct | Direct-AD2prog | Brainage64D-finetune-AD2prog |
| Direct<br>(0.6047 ± 0.0587) | | **2.90e-5** | **3.00e-4** |
| Direct-AD2prog<br>(0.7426 ± 0.0382) | | | 0.756 |
| Brainage64D-finetune-AD2prog<br>(0.7375 ± 0.0440) | | | |

Consistent with previous studies (Oh et al., 2019; Lian et al., 2020; Wen et al., 2020), our results suggest that MCI progression prediction can be improved by transferring features from previously trained AD classification models. However, we did not observe any additional benefit from leveraging features of a pretrained brain age model.

### 3.5. Even when sample size is small, lehveraging a pretrained brain age model does not improve MCI prediction over training a model directly

Adapting a pretrained model (trained from large datasets) for a new classification task might be more advantageous when the sample size available for the new task is small. Figure 8 shows the MCI prediction AUC (across 50 training-validation-test splits) for Direct-AD2prog and Brainage64D-finetune-AD2prog across different development set sizes. We did not consider the “direct” approach for this analysis, given its poor performance (Figure 7). Table 9 shows the actual AUC values and p values from comparing Direct-AD2prog and Brainage64D-finetune-AD2prog. Across all sample sizes, there was no statistical difference between Direct-AD2prog and Brainage64D-finetune-AD2prog. Overall, this suggests that even when sample size is small, there was not a significant advantage in leveraging features from a pretrained brain age model.

**Figure 8.**
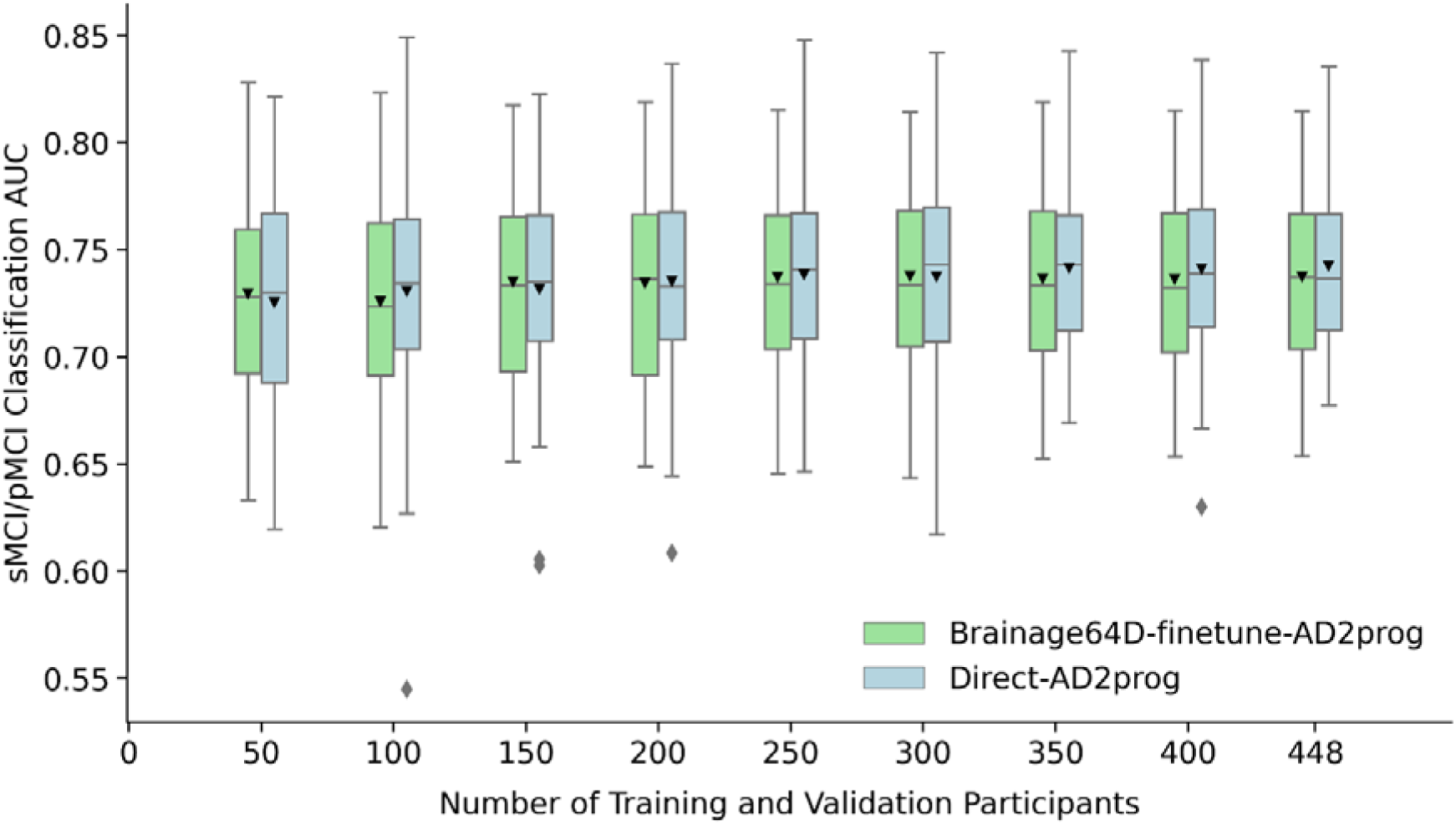
MCI progression prediction AUC of Brainage64D-finetune-AD2prog and Direct-AD2prog across different development set sizes. Boxplots showing test AUC across different development set sizes (number of training and validation participants). Test sets were identical across development set sizes, so AUCs were comparable across sample sizes. For each boxplot, the horizontal line indicates the median across 50 test AUC values. Triangle indicates the mean. The bottom and top edges of the box indicate the 25th and 75th percentiles, respectively. Outliers are defined as data points beyond 1.5 times the interquartile range. The whiskers extend to the most extreme data points not considered outliers.

**Table 9.**
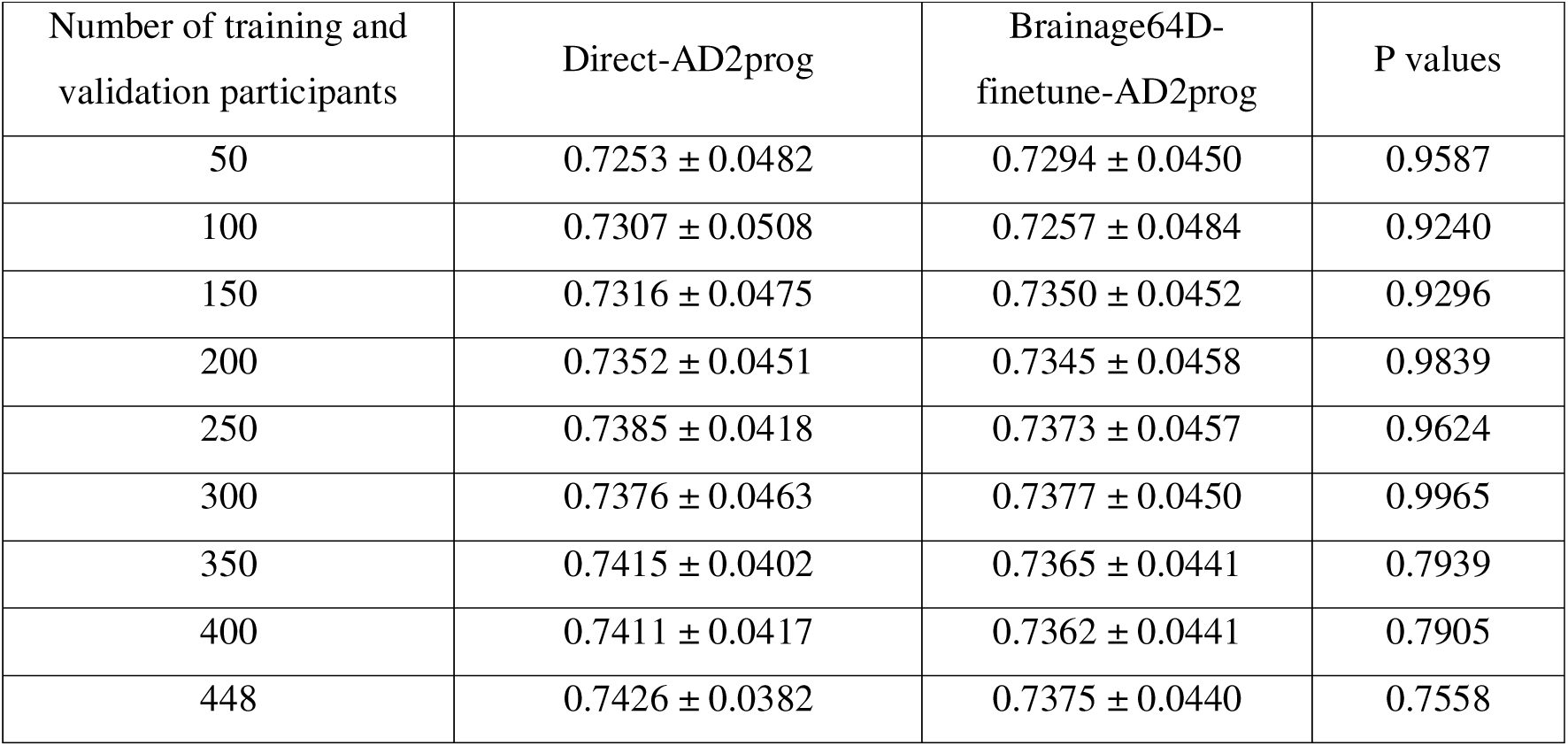
MCI prediction AUC (mean ± std) and p values comparing Direct-AD2prog and Brainage64D-finetune-AD2prog across different development set sizes (i.e., number of training and validation participants). Given its poor performance (Figure 7), we did not include the “Direct” approach in this analysis. Across all sample sizes, there was no statistically significant difference.

| Number of training and validation participants | Direct-AD2prog | Brainage64D-finetune-AD2prog | P values |
| --- | --- | --- | --- |
| 50 | 0.7253 ± 0.0482 | 0.7294 ± 0.0450 | 0.9587 |
| 100 | 0.7307 ± 0.0508 | 0.7257 ± 0.0484 | 0.9240 |
| 150 | 0.7316 ± 0.0475 | 0.7350 ± 0.0452 | 0.9296 |
| 200 | 0.7352 ± 0.0451 | 0.7345 ± 0.0458 | 0.9839 |
| 250 | 0.7385 ± 0.0418 | 0.7373 ± 0.0457 | 0.9624 |
| 300 | 0.7376 ± 0.0463 | 0.7377 ± 0.0450 | 0.9965 |
| 350 | 0.7415 ± 0.0402 | 0.7365 ± 0.0441 | 0.7939 |
| 400 | 0.7411 ± 0.0417 | 0.7362 ± 0.0441 | 0.7905 |
| 448 | 0.7426 ± 0.0382 | 0.7375 ± 0.0440 | 0.7558 |

## 4. DISCUSSION

In this study, we evaluated the utility of brain age models in generating specific markers of brain health in the domain of AD dementia. We found that models directly trained to predict AD-related outcomes (i.e., direct models) performed as well as, or even better than, brain age derived models. Even when training data was scarce (N = 50), brain age derived models did not statistically outperform direct models. Overall, given the widespread availability of brain data with age-only information, we believe that brain age can still be useful as a marker of general brain health. However, our results suggest that current large-scale brain age models do not offer a strong advantage for predicting specific health outcomes.

### 4.1. Brain age as transfer learning

To interpret our results, it is useful to think of brain age derived models as a form of transfer learning (Weiss et al., 2016; Zhuang et al., 2021), which can be broadly defined as using past experience from one or more source tasks to improve learning on a target task (Hospedales et al., 2022). In the case of brain age derived models, we first train a model to predict age in a large dataset, followed by model transfer to predict another phenotype in a new dataset. There are a few factors that might influence the success of transfer learning in the current study.

The first factor to consider is sample size. Given that the brain age model is trained from a very large dataset (N > 50,000), the hope is that the features will be more robust than those learned directly from a small dataset with target outcomes of interest (N < 1000), thus improving the performance of the target outcomes. We note that in the scenario with very small sample sizes (N = 50), there were >1000 times more data for pretraining the brain age model than the target task, yet brain age models did not yield a significant advantage over direct models.

The second factor to consider is the similarity between source and target tasks. Transfer learning is easier if source and target tasks are more similar. Conversely, transfer learning is harder if source and target tasks are more different. However, given that age is the strongest risk factor for AD dementia (Hou et al., 2019) and since the brain age gap is more sensitive to dementia compared with eight other disorders (Kaufmann et al., 2019), we believe the source and target tasks in our study are reasonably well-aligned.

The third factor is the impact of well-documented site differences which might arise from differences in MRI scanners, acquisitions, population differences and/or preprocessing differences (Fortin et al., 2018; Pomponio et al., 2020; An et al., 2024; Dular & Špiclin, 2024). These site differences may degrade model transfer. However, some studies have suggested that training models on large diverse datasets could overcome site differences without having to explicitly perform harmonization (Abraham et al., 2017; P. Chen et al., 2024), including the original study whose pretrained brain age model was adopted in the current study (Leonardsen et al., 2022).

Other studies have advocated more extensive preprocessing to alleviate inter-site factors (Dular et al., 2024) or that calibration should be considered (Smith et al., 2019; Wood et al., 2024). Following Leonardsen and colleagues (Leonardsen et al., 2022), the BAG and brainage64D models in the current study did not perform any calibration. On the other hand, transfer learning was used for the remaining brain age model approaches.

Overall, when considering all three factors, we note that it was not a foreordained result that brain age derived models and direct models would have similar performance. Indeed, an interesting finding is that BAG exhibited the worst performance, and finetuning did not improve the performance of BAG. This suggests that while BAG might be a powerful marker of general brain health, too much information is lost by reducing a person’s brain health to a single number, thus diminishing its utility for predicting specific brain health outcomes.

Furthermore, Brainage64D exhibited worse performance than Brainage64D-finetune and Direct models. On the other hand, Brainage64D-finetune and Direct models performed similarly well in the AD classification task. Brainage64-finetune-AD2prog and Direct-AD2prog models also performed similarly well in the MCI progression task. Overall, this suggests the importance of finetuning brain age models to new tasks potentially due to site differences or due to task misalignment between predicting chronological age and AD-related health outcomes.

Finally, it is important to note that brain age derived models are just one approach of transfer learning. There is a plethora of transfer learning approaches in the brain imaging (Deepak & Ameer, 2019; Malik & Bzdok, 2022; Mei et al., 2022) and machine learning (Palatucci et al., 2009; Bengio, 2011; Shin et al., 2016) literature. We believe that the idea of translating models trained from large-scale datasets to predict new phenotypes in small datasets remains a promising one. However, similarity between the source and target tasks is an important factor that needs to be taken care of to maximize transfer learning performance (He et al., 2022; P. Chen et al., 2024; Wulan et al., 2024).

Indeed, a recent study suggests that age prediction based on neuropsychological measures leads to better MCI progression prediction than BAG from brain imaging data (Garcia Condado et al., 2023). Furthermore, there are also studies that have developed biological age models based on mortality (Levine et al., 2018). It is possible that brain age models trained on mortality, rather than chronological age, could yield better markers of specific brain health. We leave this to future work.

### 4.2. Causality vs predictive utility

Age is the largest risk factor for AD dementia (van der Flier & Scheltens, 2005; Daviglus et al., 2010). The risk of AD increases exponentially after 65 years old. The prevalence of AD dementia is ∼50% for individuals ≥ 95 years old in the United States (Hou et al., 2019). A seminal review described nine cellular and molecular hallmarks of aging (López-Otín et al., 2013), all of which have been implicated in AD (Hou et al., 2019). Furthermore, using the same pretrained brain age model from the current study, AD risk genes have been shown to have a causal effect on brain age gap (Leonardsen et al., 2023). Together, these results suggest a potential causal link between AD and brain age.

However, it is important to note that a causal link is not sufficient to make chronological age a good pretraining target. For example, lead exposure causally reduces Intelligence Quotient (IQ), but the effect size is modest (r = 0.12; Meyer et al., 2001). Therefore, using lead exposure alone to predict IQ would probably lead to poor prediction performance. Similarly, even if brain aging causally contributes to AD, its predictive signal, particularly when compressed into a single scalar (brain age gap), can be weak. Therefore, our findings should not be taken as evidence against a causal link between brain aging and AD.

Indeed, the goal of the current study is not to evaluate the causal link between AD and brain age, but to evaluate chronological age prediction as a pretraining task. The assumption that chronological age prediction is a good pretraining task is implicit or explicit in many brain age studies (Kaufmann et al., 2019; Bashyam et al., 2020; Lee et al., 2022; Leonardsen et al., 2022; Moguilner et al., 2024). As already mentioned in the introduction, among nine brain disorders, brain age gap was the most sensitive to dementia (Kaufmann et al., 2019), suggesting that AD classification is a natural benchmark.

In the case of MCI progression prediction, it is common to train models on CN vs AD classification and transfer the models (or their learned features) to predict MCI progression (Liu et al., 2018; Bron et al., 2021; Kang et al., 2023). This practice assumes that features distinguishing CN from AD are informative about disease trajectory, consistent with our results (Figure 7). Therefore, given that brain age is especially sensitive to dementia (Kaufmann et al., 2019), MCI progression prediction is also a natural testbed for evaluating brain age models.

### 4.3. Alzheimer’s disease vs psychiatric disorders

Due to pronounced atrophy patterns in AD, our tasks of AD classification and MCI progression prediction using T1 MRI are probably easier than psychiatric symptoms or disorder prediction using noisy resting-state functional connectivity (Chen et al., 2022; Traut et al., 2022; Winter et al., 2024). Extending our framework to alternative imaging modalities and to psychiatric conditions is a promising direction for future research.

We also note that MCI progression prediction represents a substantially more challenging task than AD classification, as reflected in the lower overall performance across models. However, the relative performance between brain age and direct models remained comparable. This suggests that while prediction accuracy may drop for both approaches in the context of psychiatric disorders and noisier modalities like resting-state functional MRI, such a setting would not necessarily confer an advantage to brain age models over direct models.

Furthermore, from a theoretical perspective, chronological age is a biologically meaningful pretraining target in AD, where aging is the primary risk factor (Section 4.2). In contrast, psychiatric disorders often emerge during adolescence or early adulthood, a developmental window characterized by rapid brain maturation and heightened plasticity, rather than as a function of aging per se (Paus et al., 2008). As such, the biological relevance of age as a pretraining target may be weaker in the context of psychiatric disorders.

Empirical evidence supports this distinction. Kaufmann et al. (2019) found that brain age gap based on T1-weighted MRI was markedly more sensitive to dementia (Cohen’s d = 1.03) than to psychiatric disorders such as major depressive disorder (d = 0.10), bipolar disorder (d = 0.29), schizophrenia (d = 0.51), ADHD (d = 0.29), or autism (d = 0.07). This further suggests that brain age models may be even less well suited for psychiatric disorder or symptom prediction than for AD classification or MCI progression prediction.

### 4.4. Limitations & future work

A limitation of the current study is that we only considered one pretrained brain age model (Leonardsen et al., 2022). We believe that this model remains the best (or one of the best) in the field, but we do not preclude the possibility that other brain age models trained on even larger and more diverse datasets might yield better transfer learning results. However, new brain age models are typically trained to improve chronological age prediction (Dartora et al., 2024; Kalc et al., 2024). In doing so, the models might learn better features for predicting age, but not necessarily for predicting other phenotypes.

Another limitation is that since the pretrained brain age model used the SFCN architecture (Leonardsen et al., 2022), the direct models also had to use the same architecture for the comparison to be fair. We note that the SFCN architecture was used by the winner of the Predictive Analysis Challenge 2019 for brain age prediction (Peng et al., 2021). A recent study also demonstrated using the SFCN architecture that pretraining with a comprehensive set of targets (spanning multiple phenotypic domains) could improve the prediction of new phenotypes in a small dataset over direct models (Wulan et al., 2024). Overall, we believe that the SFCN architecture does not unfairly advantage direct models over brain age models in our comparison.

However, what is less obvious is whether using a different neural architecture might improve direct models. Studies in the past five years have reported AD classification AUC ranging from 0.82 to 0.99 (Bae et al., 2020; Bashyam et al., 2020; Qiu et al., 2020; Ocasio & Duong, 2021; Ashtari-Majlan et al., 2022; Khatri & Kwon, 2022; Leonardsen et al., 2022; Lu et al., 2022; Zheng et al., 2022; Kang et al., 2023; Aghdam et al., 2024; Q. Chen et al., 2024; Khatri & Kwon, 2024; Kumari et al., 2024; Yin et al., 2024; Zarei et al., 2024; Zhu et al., 2024), and MCI progression prediction AUC ranging from 0.62 to 0.99 (Gao et al., 2020; Lian et al., 2020; Nanni et al., 2020; Bron et al., 2021; Li et al., 2021; Ocasio & Duong, 2021; Ashtari-Majlan et al., 2022; Lu et al., 2022; Garcia Condado et al., 2023; Kang et al., 2023; Luo et al., 2024; Zhou et al., 2024; Li et al., 2025; Liu et al., 2025). Therefore, while our AD classification AUC (0.92) and MCI progression prediction AUC (0.74) are well within the range of these previous studies.

The better performance reported by some of the previous studies might be due to several reasons. First, some studies used additional information beyond a single T1 in the current study, e.g., neuropsychological scores (Qiu et al., 2020; Garcia Condado et al., 2023; Luo et al., 2024), fMRI (Khatri & Kwon, 2022; Liu et al., 2025) and other structural imaging modalities (Zhu et al., 2024; Li et al., 2025). Second, following Leonardsen and colleagues (2022), our study performed a careful matching of covariates (age, sex and scanner) between AD and CN, as well as between sMCI and pMCI. Furthermore, we only considered one T1 per participant during training and validation. As a result, a number of studies reporting higher prediction performance utilized much larger training and validation sample sizes (Lu et al., 2022; Q. Chen et al., 2024; Kumari et al., 2024). Finally, a third reason could be better neural architecture or algorithm, e.g., vision transformer (Lian et al., 2020; Aghdam et al., 2024; Khatri & Kwon, 2024) or different pretraining procedure (Bae et al., 2020; Kang et al., 2023).

Our study aims to evaluate the effectiveness of using chronological age as a pretraining target, a common yet often implicit assumption in brain age research, as opposed to optimizing neural network architectures or pretraining strategies for AD classification or MCI progression prediction. Direct comparisons across studies are limited by variations in datasets, cohort characteristics, and problem formulations. However, Leonardsen et al. (2022) offers a meaningful point of reference, as we closely replicated their experimental setup. Although their study did not include MCI progression prediction, the similarity in AD classification performance between their model (AUC = 0.83) and our Brainage64D model (AUC = 0.84) supports the validity of our brain age implementation.

## 5. CONCLUSION

Brain age is a powerful marker of *general* brain health. However, even with a brain age model pretrained on a large number of participants, the resulting prediction of *specific* brain health outcomes was not better than models directly trained to predict the same outcomes. Overall, our results suggest that chronological age prediction might not be a strong pretraining target for transfer learning to predict specific health outcomes.

## ACKNOWLEDGEMENTS

This research is supported by the NUS Yong Loo Lin School of Medicine (NUHSRO/2020/124/TMR/LOA), the Singapore National Medical Research Council (NMRC) LCG (OFLCG19May-0035), NMRC CTG-IIT (CTGIIT23jan-0001), NMRC OF-IRG (OFIRG24jan-0006; OFIRG24jul-0049), NMRC STaR (STaR20nov-0003), Singapore Ministry of Health (MOH) Centre Grant (CG21APR1009), and the United States National Institutes of Health (R01MH133334). Any opinions, findings and conclusions or recommendations expressed in this material are those of the authors and do not reflect the views of the funders. Data collection and sharing for this project was funded by the Alzheimer’s Disease Neuroimaging Initiative (ADNI) (National Institutes of Health Grant U01 AG024904) and DOD ADNI (Department of Defense award number W81XWH-12-2-0012). ADNI is funded by the National Institute on Aging, the National Institute of Biomedical Imaging and Bioengineering, and through generous contributions from the following: AbbVie, Alzheimer’s Association; Alzheimer’s Drug Discovery Foundation; Araclon Biotech; BioClinica, Inc.; Biogen; Bristol-Myers Squibb Company; CereSpir, Inc.; Cogstate; Eisai Inc.; Elan Pharmaceuticals, Inc.; Eli Lilly and Company; EuroImmun; F. Hoffmann-La Roche Ltd and its affiliated company Genentech, Inc.; Fujirebio; GE Healthcare; IXICO Ltd.;Janssen Alzheimer Immunotherapy Research & Development, LLC.; Johnson & Johnson Pharmaceutical Research & Development LLC.; Lumosity; Lundbeck; Merck & Co., Inc.;Meso Scale Diagnostics, LLC.; NeuroRx Research; Neurotrack Technologies; Novartis Pharmaceuticals Corporation; Pfizer Inc.; Piramal Imaging; Servier; Takeda Pharmaceutical Company; and Transition Therapeutics. The Canadian Institutes of Health Research is providing funds to support ADNI clinical sites in Canada. Private sector contributions are facilitated by the Foundation for the National Institutes of Health (www.fnih.org). The grantee organization is the Northern California Institute for Research and Education, and the study is coordinated by the Alzheimer’s Therapeutic Research Institute at the University of Southern California. ADNI data are disseminated by the Laboratory for Neuro Imaging at the University of Southern California.

## DATA AVAILABILITY STATEMENT

This study utilized publicly available data from ADNI (https://adni.loni.usc.edu/), AIBL (https://aibl.org.au/), and MACC (https://medicine.nus.edu.sg/macc-2/) datasets. Data can be accessed via data use agreements.

